# Diagnosis of planktonic trophic network dynamics with sharp qualitative changes

**DOI:** 10.1101/2023.06.29.547055

**Authors:** Cedric Gaucherel, Stolian Fayolle, Raphael Savelli, Olivier Philippine, Franck Pommereau, Christine Dupuy

## Abstract

Trophic interaction networks are notoriously difficult to understand and to diagnose (i.e., to identify contrasted network functioning regimes). Such ecological networks have many direct and indirect connections between species, and these connections are not static but often vary over time. These topological changes, as opposed to a dynamic on a static (frozen) network, can be triggered by natural forcings (e.g., seasons) and/or by human influences (e.g., nutrient or pollution inputs). Aquatic trophic networks are especially dynamic and versatile, thus suggesting new approaches for identifying network structures and functioning in a comprehensive manner.

In this study, a qualitative model was devised for this purpose. Applying discrete-event models from theoretical computer science, a mechanistic and qualitative model was developed that allowed computation of the exhaustive dynamics of a given trophic network and its environment. Once the model definition is assumed, it provides all possible trajectories of the network from a chosen initial state. In a rigorous and analytical approach, for the first time, we validated the model on one theoretical and two observed trajectories recorded at freshwater stations in the La Rochelle region (Western France). The model appears to be easy to build and intuitive, and it provides additional relevant trajectories to the expert community. We hope this formal approach will open a new avenue in identifying and predicting trophic (and non-trophic) ecological networks.

## INTRODUCTION

Trophic networks (TNs) form the backbone of ecosystem functioning, as they simultaneously condition food acquisition, prey and predator demography, individual and population behaviors, and phenotype selection, among other consequences (Lindeman 1942, Johnson 2000, Majdi et al. 2018). Trophic processes are responsible for most matter and energy fluxes within ecosystems, but the fates and properties of ecosystems are hard to predict, mainly due to the lack of knowledge (Mouquet et al. 2015). Trophic processes have been extensively studied in ecology, although mostly considered as frozen in time, i.e., with a fixed (or static) topology and fixed signed interactions. This simplification may be due to a lack of long-term data as well as to equation-based models dedicated to flux and abundance variations on a frozen network (e.g., Thébault and Fontaine 2010, Kéfi et al. 2015). In this study, we provide an original framework to handle TNs with sharply changing structures and to model their possible dynamics.

To date, TNs have been difficult to understand and handle, in other words, hard to *diagnose* between contrasted functioning under changing environmental conditions. Any new functioning involves specific ecosystem components and interactions, thus justifying why we have focused on qualitative functioning *regimes* rather than quantitative component abundances and interaction intensities. In addition, TNs usually gather a large number of populations or species in an even larger number of trophic interactions. Hence, understanding trophic dynamics would require not only modeling a large and realistic number of components but also being able to calibrate the weights (coefficients) of each component and each interaction involved (Ings et al. 2009, Wallach et al. 2017, Majdi et al. 2018). For this reason, most trophic models to date have focused on wide categories of populations, with functional categories, such as carnivores, herbivores, and/or detritivores (e.g., Thébault and Fontaine 2010), and approximate their trophic parameters. Even powerful models aimed at bypassing such limitations, such as qualitative models based on differential equation systems, are limited in size (May 1973, Dambacher et al. 2003).

There is an even more pronounced limitation of trophic studies in ecology, as they mostly assume a frozen network of interaction (Thébault and Fontaine 2010, Kéfi et al. 2016). Not only is it harder to handle a network that changes in terms of topology (structure), but it is also not known how such a network may change over time and, thus, how to model it. Indeed, as soon as the study covers several generations of some of the species involved in the network, other species may invade and/or become extinct (Mooney and Hobbs 2001, Warren et al. 2005). Hence, these events greatly modify the network structure and, in turn, the system dynamics. Equation-based models are not well suited to handle dynamic systems *on* dynamic structures (sometimes called DS^2^, Giavitto and Michel 2003), whereas certain tools developed in theoretical computer science are perfectly adapted to this task. In particular, discrete-event systems such as graph transformations or Petri nets are able to handle sharply changing networks by formalizing the way components and interactions can appear or disappear (König et al. 2018, Gaucherel and Pommereau 2019). While graph transformations directly add/remove some nodes and edges, Petri nets only mimic such addition/removal by marking the presence/absence of the handled nodes and edges with some tokens. In this study, we developed a Petri net to model any interaction network topological change, and we illustrate its use on a realistic planktonic TN.

Planktonic TN models are usually composed of a fixed number of functional nodes that gather groups of individuals sharing the same ecological function. Mass fluxes (usually in carbon or nitrogen) between nodes are predefined according to trophic interactions. In a context of an emerging biological oceanography discipline and considering the limited computing resources, the first planktonic TN (or food web) models simply consisted of mass fluxes between nutrients, phytoplankton, and zooplankton nodes (Steele 1958, 1974). These so-called NPZ models (NPZD, NPZDB, or even NPZF when detritus, bacteria, or fishes are comprised, respectively) coupled to observed or simulated physical conditions have demonstrated their predictive ability to capture bulk system properties (biomass and primary production) at both regional and global scales (Mitra et al. 2007, Kriest et al. 2010, Oke et al. 2013, Hernández-Carrasco et al. 2014, Turner et al. 2014, Kumar and Kumari 2015).

To better understand biogeochemical cycling (e.g., export fluxes, carbon sequestration, organic matter recycling, microbial loop), planktonic TN should be delineated, and planktonic compartments in models could thus be refined into Planktonic Functional Types (PFTs). Plankton groups are thus defined according to common ecological functions (e.g., nitrogen fixers, calcifiers, and silicifiers), sizes (e.g., picophytoplankton, nanophytoplankton, microphytoplankton), and/or key taxonomic groups (e.g., diatoms, flagellates) (Le Fouest et al. 2013, Villaescusa et al. 2016, Kerimoglu et al. 2017, Petersen et al. 2017, Maar et al. 2018, Meddeb et al. 2019). However, refinements of planktonic TN models greatly complicate model formalization and parametrization, as well as requiring more data, which increases uncertainties in terms of the model outcomes and fluxes between defined groups (Anderson 2005).

To address gaps in ecological knowledge and lacking data, inverse modeling is aimed at deriving flows of energy within TNs from simple biomass estimates and rate measurements. Vézina and Platt (1988) were the first to use this for inferring mass fluxes through a planktonic TN in the English Channel. Inverse modeling is, therefore, advantageous when dealing with underdetermined systems and results in a space of possible solutions that fulfill a set of linear equalities and inequalities. A preferred solution is then selected by optimization or statistical methods. While vital rates and biomass can be readily measured for high trophic levels (e.g., fishes), their quantification for low trophic levels (e.g., bacteria, autotrophic plankton) remains uncertain and questions the robustness of inverse modeling for the study of planktonic compartments (Vernet et al. 2017, Saint-Béat et al. 2018). Overall, biological constants (production, consumption, assimilation), biomass, and ecological interactions are, therefore, not easy to measure in planktonic TNs, resulting in an oversimplification of planktonic TN models (Anderson 2005, Flynn 2006). For all of these reasons, our main objective in this study consisted of developing a model able to identify (diagnose) any qualitative functioning regimes of the same TN under changing environmental conditions.

We here addressed the leading question: what are all the possible trajectories (pathways) of such an aquatic TN? A trajectory is defined here as a sequence of TN states (regimes) and transitions in time, possibly exhibiting bifurcations and not necessarily being quantitative. More specifically, we aimed to identify the various qualitative regimes the system can reach between winter and summer environmental conditions. As a second subquestion, we assessed whether a detailed model could exhibit new or counter-intuitive TN trajectories. We assumed that the system may be qualified and thus exhibit a finite number of states, computed and gathered into a Petri net *state space* (Pommereau 2010, Reisig 2013). A detailed and automatic analysis of this state space then exhaustively provides the possible fates (e.g., stabilities, collapses, if any) of the studied system. We chose to illustrate this original method with a well-studied plankton TN in wetlands, namely freshwater marshes of the Charente Maritime region (Western France, Tortajada et al. 2011). Such a system is well instrumented (measured) and will provide an expected theoretical trajectory of changing TN, as well as two observed trajectories at different stations (Masclaux et al. 2014). The succession of planktonic TN and the different regimes of the planktonic TN are well known according to the season (Masclaux et al. 2014). We developed the corresponding Petri net of this system and then validated it, for the first time, on theoretical and observed trajectories. We finally discuss the power and drawbacks of such discrete and qualitative models for trophic ecology.

## MATERIALS AND METHODS

### Aquatic trophic networks

The Charente-Maritime marshes of the French Atlantic coast (Fig. 1) are the second largest French wetland zone (more than 1000 km^2^). The type of freshwater marshes is unreplenished drained marshes, which constitute a significant artificial hydrographic network of channels and ditches. To mitigate and prevent drying of the marshes, locks control the channels and ditches (for more details, see Masclaux et al. 2014). Samples of the planktonic TN were recorded at two stations (stations A and B) on successive dates (eight weeks during winter and spring 2012) to reconstruct the network trajectories and their environment over time (Masclaux et al. 2014). All of the details have been presented in the publication by Masclaux et al. (2014). Briefly, the water parameters studied were the temperature, nutrients (nitrates, etc.), the dissolved organic matter (*DOC*) concentration, the biomass and production of bacteria, the biomass and primary production of phytoplankton by size class (microphytoplankton for > 20 μm; nanophytoplankton for 3–20 μm; and picophytoplankton for < 3 μm), the bacterial biomass, the protozoa biomass, and the metazoan microzooplankton and mesozooplankton biomasses. Different fluxes between preys and predators were measured: micro- and mesozooplankton grazing rates on the three phytoplankton size classes, as well as mesozooplankton grazing rates on protozoa (Masclaux et al. (2014). The TN regimes were determined with a hierarchical ascendancy classification (HAC, Euclidean distance, and Ward method). The planktonic TN regime changed during the winter to spring transition, from biological winters, followed by herbivorous TNs, to finally reach TNs qualified as multivorous and distinguishing three levels of multivory (weakly multivorous, multivorous, and highly multivorous) (Masclaux et al. 2014).

**Figure 1.**
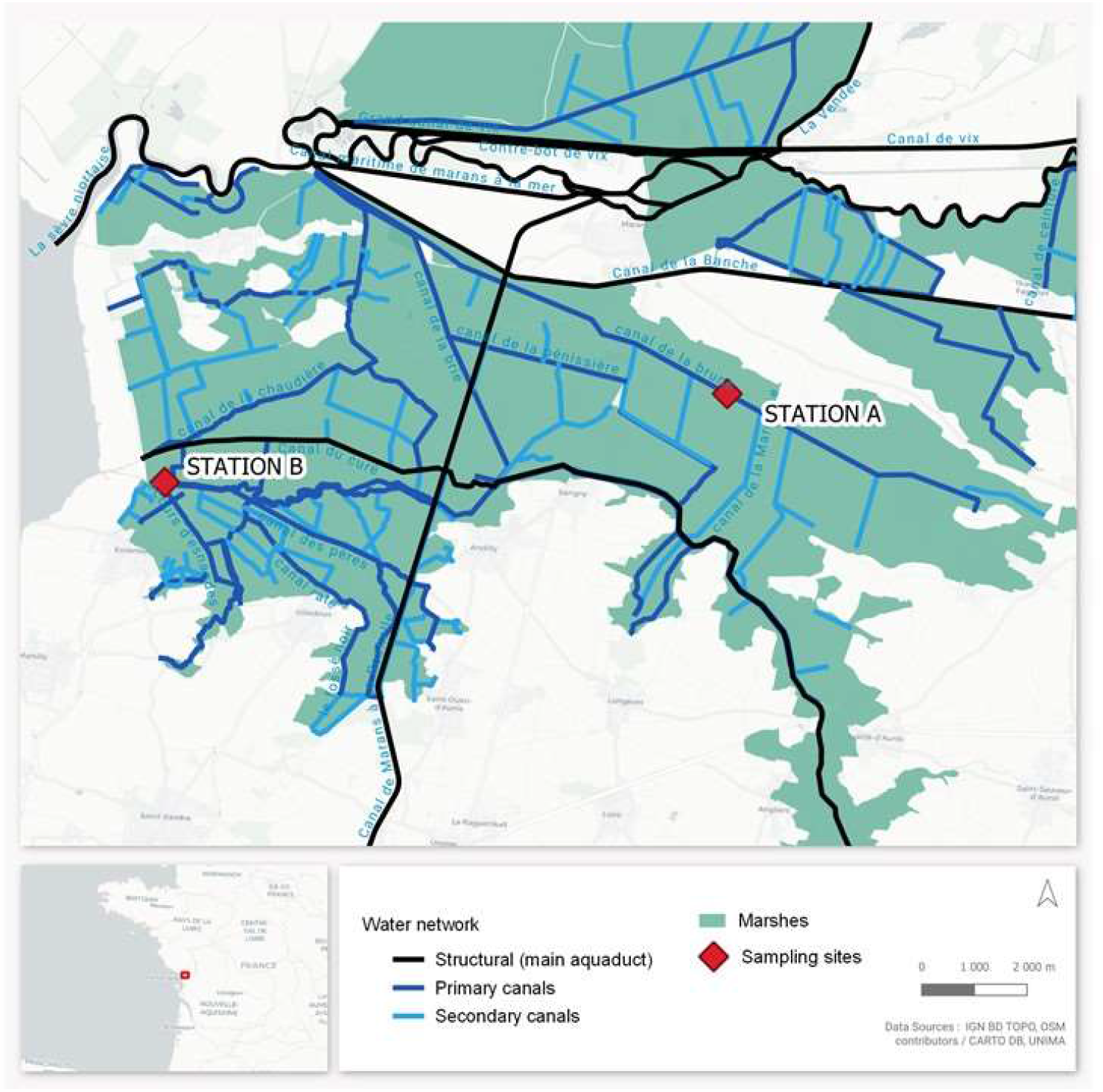
Location of the study site and the two sampled stations (A and B, inset) along the Atlantic coast of France.

The model was built with several categories of variables (Fig. 2): Phytoplankton, Zooplankton, Resources, Microbes, and Abiotic components characterizing the environment. The main functional groups were: 1) phytoplankton by size class (microphytoplankton for > 20 μm: *MicroP*; nanophytoplankton for 3–20 μm: *NanoP;* and picophytoplankton for < 3 μm: *PicoP*) all in green (Fig. 2); 2) metazoan microzooplankton (*MicroZ*) and mesozooplankton (*MesoZ*) in red; 3) resources as nitrates (*Nit*) and Dissolved Organic Matter (*DOC*) in brown; 4) microbes such as bacteria (*Bact*) and protozoa (*Proto*) in blue; and 5) abiotic variables in grey with the component *envir*, which corresponds to the temperature and light conditions and *renew* which corresponds to the possible renewal of water (i.e., flush) in the marshes depending on the rainfall and water usages (agriculture, breeding, etc.). In more detail, the planktonic TN and all the possible fluxes (interactions) between components concern grazing fluxes with some preferential predation, and potential competitions between organisms (Fig. 2). Protozoa graze on bacteria, *PicoP* and *NanoP*, and are grazed by *MicroZ* and *MesoZ. MicroZ* graze on bacteria, *PicoP, NanoP*, and *Proto. MicroP* is grazed mainly by *MesoZ*, which used *MicroZ, NanoP*, and *Proto* as preys. Two preferential interactions force the model: i) competition between bacteria and *PicoP*, suggesting that each may survive and be detrimental to the second one, and ii) preferential grazing of *MicroP* by *MesoZ* and preferential grazing of *NanoP* by *MicroZ*. In brief, plain upward edges correspond to prey-predator interactions, while dashed downward edges are the resulting detritus (toward the *DOC* variable).

**Figure 2.**
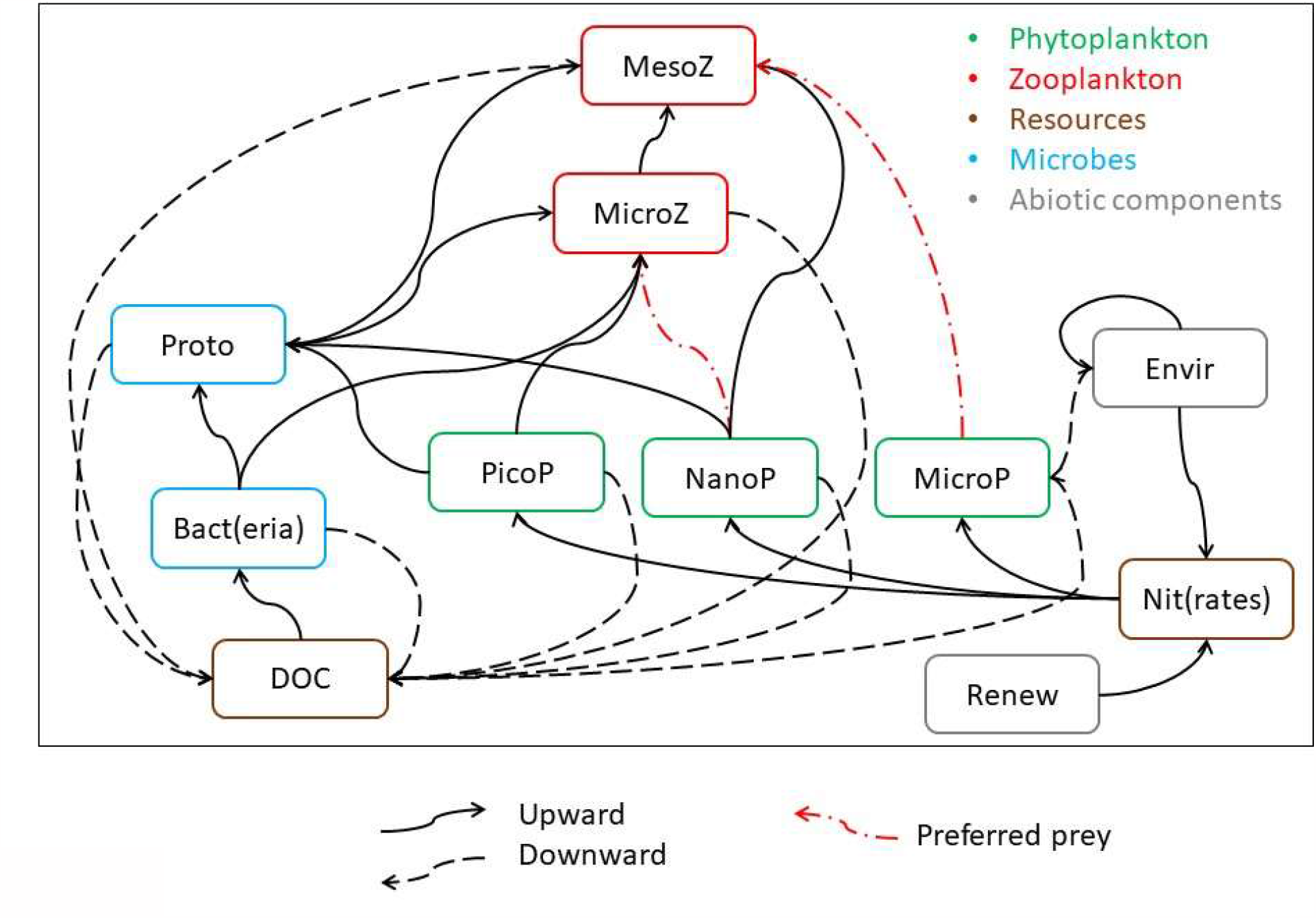
The detailed interaction network used in this study. The trophic and non-trophic components are displayed as nodes with various colors (Phytoplankton: green, Zooplankton: red, Resources: brown, Microbes: blue, and Abiotic components characterizing the environment: grey). Predation interactions are displayed as plain (upward for trophic) and dashed (downward for degradation) edges, with preferred prey populations highlighted with red doted-dashed edges.

### Petri Nets and a simplistic Prey-Predator model

We first summarize the successive steps required to build any ecosystem Petri net and we then illustrate these steps with a toy model. Our generic approach to model ecosystems has been called the EDEN (Ecological Discrete-Event Network) modeling framework and is specifically applied here to an aquatic trophic TN. Any ecosystem Petri net is developed in three successive steps (Fig. 3): i) an intuitive graph (i.e., a set of nodes and edges) is built to represent the studied ecosystem with its components and their related processes, focusing on the leading question addressed by the model (Fig. 3a); ii) this ecosystemic graph, now called the *interaction network*, is then transformed into a formal model based on a discrete-event Petri net and its associated rules (as explained in the next paragraph, Fig. 3b); iii) the Petri net is run (computed, Fig. 3c) and analyzed (Fig. 3d) to determine the entire range of the ecosystem dynamics. However, the Petri net (steps i and ii) is hidden from the (ecologist) user and is automatically built (in Python language, see Suppl. Mat.) once the ecosystem components and processes have been chosen by the ecologists. Additional technical details regarding the principle and use of Petri nets, in particular the way they are computed, can be found in the literature (Pommereau 2010, Reisig 2013, Gaucherel and Pommereau 2019).

**Figure 3.**
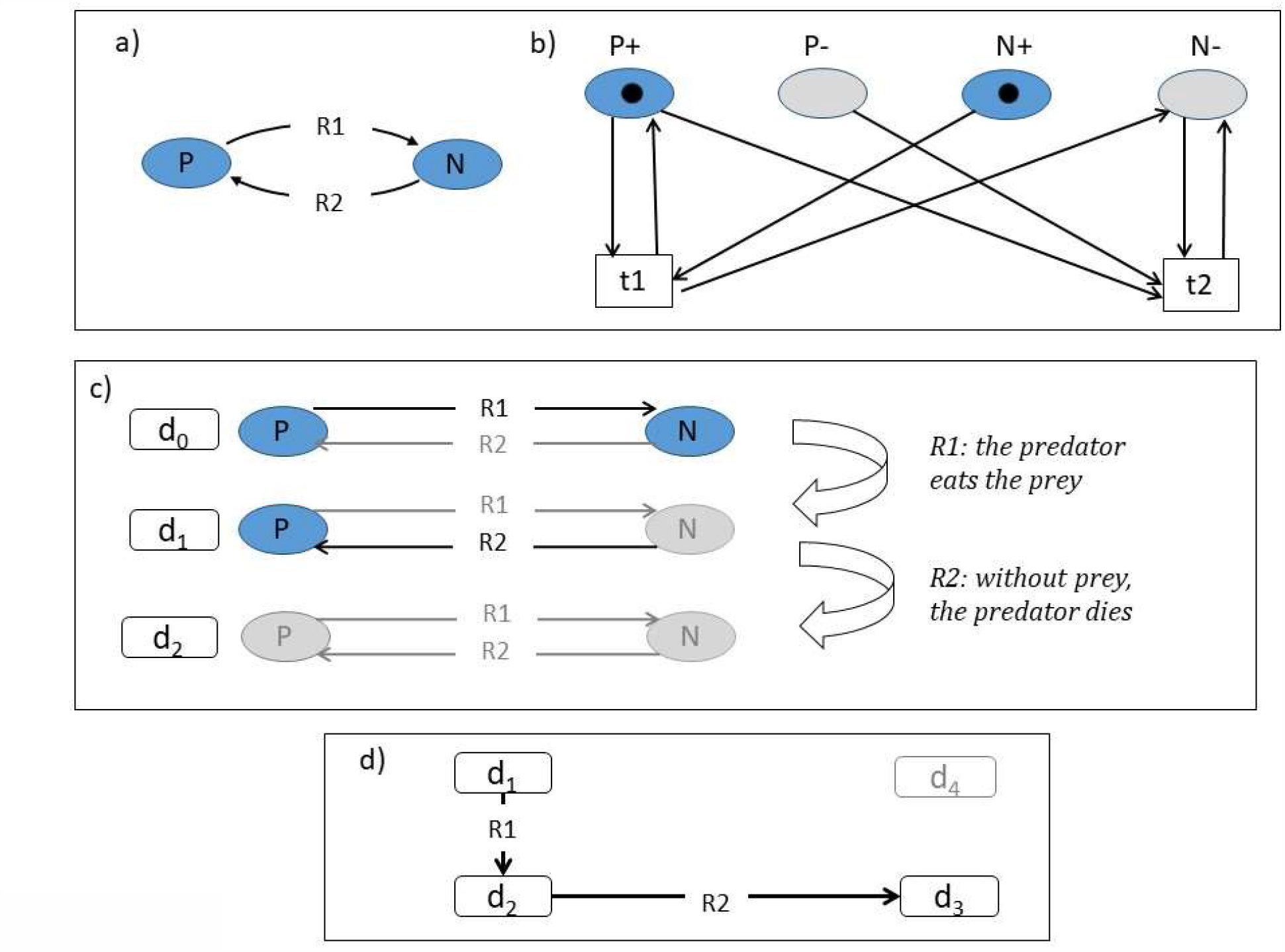
Illustration of a simplistic prey-predator system (a), with its associated Petri net (b), its qualitative dynamics (c), and the computed marking graph, also called state space (d). The system comprises two components, the prey (N) and predator (P) populations, and two interactions connecting them (rules R1 and R2, (a)). The corresponding Petri net comprises four places (P+, P-, N+, and N-) and two transitions, R1 and R2, linked by unlabeled and unweighted arcs (b). Starting with the presence of both populations, it is possible to list all system states encountered {d_1_, d_2_, and d_3_} (c), and to connect them with the rules (absent nodes and inactivated rules are displayed in grey). The net is depicted in the initial state (c), and the successive states can be deduced from the token (black dot in (a)) movements between places (b). The marking graph of the Petri net (d) is depicted with each state number {d_0_, d_1_, and d_2_} referring to the dynamics described above (b). It should be noted that a specific state of the system {d_4_} may not be reached from this initial condition and with these rules (d).

We here illustrate the basic functioning of the model using a simplistic prey-predator system (Fig. 3). Any ecosystem network can be represented as a directed graph (with parallel edges). In this graph, every material component of the ecosystem (e.g., abiotic: temperature; biotic: a population; anthropogenic: nitrate inputs) is represented by a node (or variable), with two Boolean states: “present” (the component is functionally present in the system, or above a chosen threshold, also denoted “+”) and “absent” (functionally absent from the system, or below the same threshold, denoted “-”). In the prey-predator toy model, only two nodes are defined: the prey and the predator populations. Any state of the ecosystem is defined by the set of “+” and “-” nodes (Fig. 3b), and “±” in the tables of this paper, when they may oscillate between successive states. The state of a node depends on the nodes to which it is connected, while a connection between nodes is assumed as soon as one process explicitly connects the different components (Figs. 3a-b). The rules correspond to any physicochemical and/or bio-ecological processes (e.g., if the prey population is below a chosen threshold (-), the predator population also ends up below its associated threshold), and thus represent all possible interactions between the components comprising the studied ecosystem. In the prey-predator system, only two rules are defined: R1, the predation itself: the predator eats the prey, and R2, the mortality: without prey, the predator dies (Figs. 3a and c). In the Petri net language, nodes are called places, and rules are called transitions, both being connected through oriented arcs (Fig. 3b).

### Discrete and qualitative dynamics

Any rule of such discrete-event models combines the left-hand *condition* and right-hand *realization* as: “transition’s name: condition >> realization”. For a rule to be applied, the state of the node (variable) must satisfy its application condition; the rule is then designated as *enabled*. If so, the application of the rule modifies the state of the node as stipulated in its realization part; the rule is then *fired* (i.e., executed or applied). In the prey-predator system (Fig. 3), the rules are written as R1: P+, N+ >> N- and R2: N-, and P+ >> P-. Since the rules modify node states, they change the overall system state (i.e., the state of the system aggregates all node states). Therefore, the entire system shifts from one state to another through the successive applications of the enabled rules (Fig. 3c). Computation of the defined Petri net produces the *state space*, which provides the set of all system states reachable by the rules defined (Fig. 3d). As a corollary, the system states are also connected to each other by some of these rules in the state space. The size of this state space is usually much smaller than the number of possible system states (2^n^, where n is the number of components or nodes/variables), because the computation starts from a specific initial condition and because the rules have specific application conditions. Following the computer science community, we developed certain tools to automatically divide large state spaces into *merged* (simplified) state spaces, as explained in the next subsection.

Firing a rule independently of some others often leads to unrealistic trajectories (e.g., flushing water without removing the plankton in it). Therefore, we defined *constraints*, which prevent the model from simulating such unrealistic trajectories. Constraints have a condition and a realization part, just as rules *stricto sensu* do, and the model inevitable (mandatory) transitions given the system state. The sole difference between rules and constraints is that constraints have priority over stricto sensu rules. In the prey-predator system, the system state d_1_ = {N-, P+} is unrealistic; so, the rule R2 has to be transformed into a constraint C1: N-, P+ >> P-. From a given state, the model first simulates all trajectories opened up by the defined constraints, and then, when all the system states obtained are realistic (i.e., there is no longer any enabled constraint), only the enabled rules are fired (Fig. 3d). It should be noted that the modeled system can remain an indefinite time in any of the computed states, as no rule forces it to leave the qualitative state (i.e., the system can experience quantitative dynamics, yet without sharp qualitative changes). In brief, the discrete model proposed here is qualitative, mechanistic (the processes are explicit), non-deterministic (no stochasticity yet several possible outcomes from each state), and asynchronous (i.e., all rules are applied as soon as possible, no rule conflict) (Reisig 2013, Gaucherel and Pommereau 2019). The EDEN models are also causal and chronological yet non-temporized, i.e., transitions and time steps are not quantified (and not probabilized).

### TN trajectories and validation methodology

The theoretical plankton TN modeled here combines nine different components, associated with the dominant functional groups that may be present in the channel freshwater marshes, and two additional components featuring environmental conditions (Table 1, Fig. 2). To connect them, we defined 34 processes and seven constraints encompassing at least four trophic levels (Tables 2-3, Fig. 2). To validate the TN model, one theoretical trajectory was defined and two observed trajectories were recorded at two distant stations (Supplementary Materials, Tables SM1-2). For the model to be validated, we expect not only to detect these successive states (e.g., {S0, S1, S2, S3}) in the modeled state space but also to detect them in the correct succession order. To determine whether the model was able to recover the expected trophic regimes, we tested two variants of the model: i) the full model intending to encompass the TN functioning, and ii) a similar model (called *seasonal*) yet without a return to winter conditions (R0, Table 2), thus resulting in the model being stuck in summer conditions. The model starts in winter conditions or with a flush in summer conditions, with only the node *Renew* present, which returns a source of inorganic nitrogen to the system (Table 1).

**Table 1.** The plankton TN components and their associated modeled variables, with their abbreviations and descriptions. Whether these ecological components are present (+) or absent (-) in the initial state is also indicated (second column).

| Acronym | In initial state | Description |
| --- | --- | --- |
| Bact | - | Bacteria |
| PicoP | - | Picophytoplankton |
| NanoP | - | Nanophytoplankton |
| MicrP | - | Microphytoplankton |
| Proto | - | Protozoa |
| MicrZ | - | Microzooplankton (metazoa) |
| MesoZ | - | Mesozooplankton |
| DOC | - | Dissolved organic matter |
| Nit | - | Nitrates, linked to seasons or flush |
| Envir | - | Environmental and climate conditions |
| Renew | + | Water and Nitrate inputs (rain or humans) |

**Table 2.**
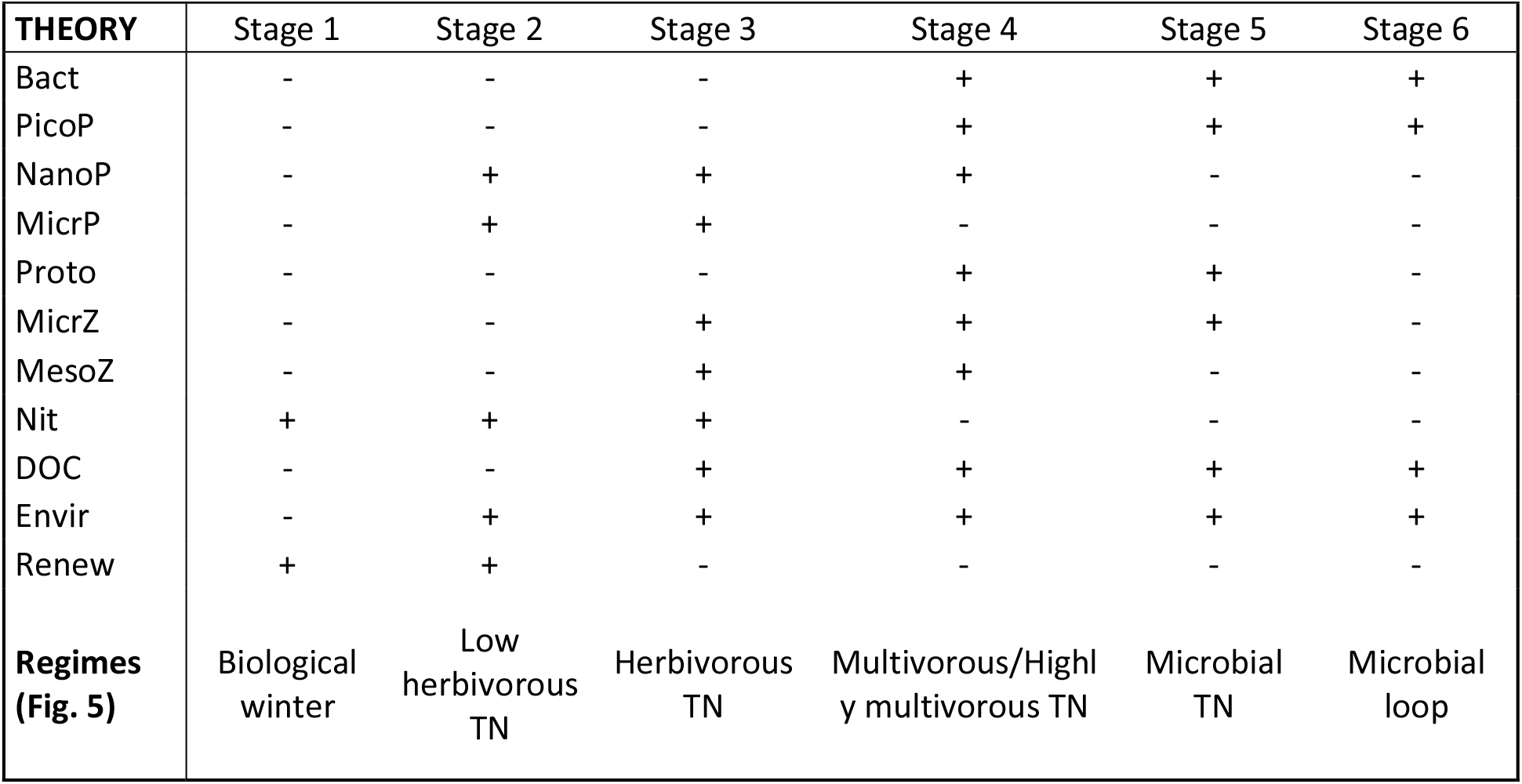
Trajectories of the theoretical expert elicitation and observed at stations A and B. For each trajectory, theoretical and observed regimes are listed in columns, and present (+)/absent (-) components of the trophic network are listed in rows. The corresponding regimes displayed in Figs. 5a-c are listed in the last row of each trajectory, with a single index A1 to A3 and B1 to B4 for successive regimes.

**Table 3.**
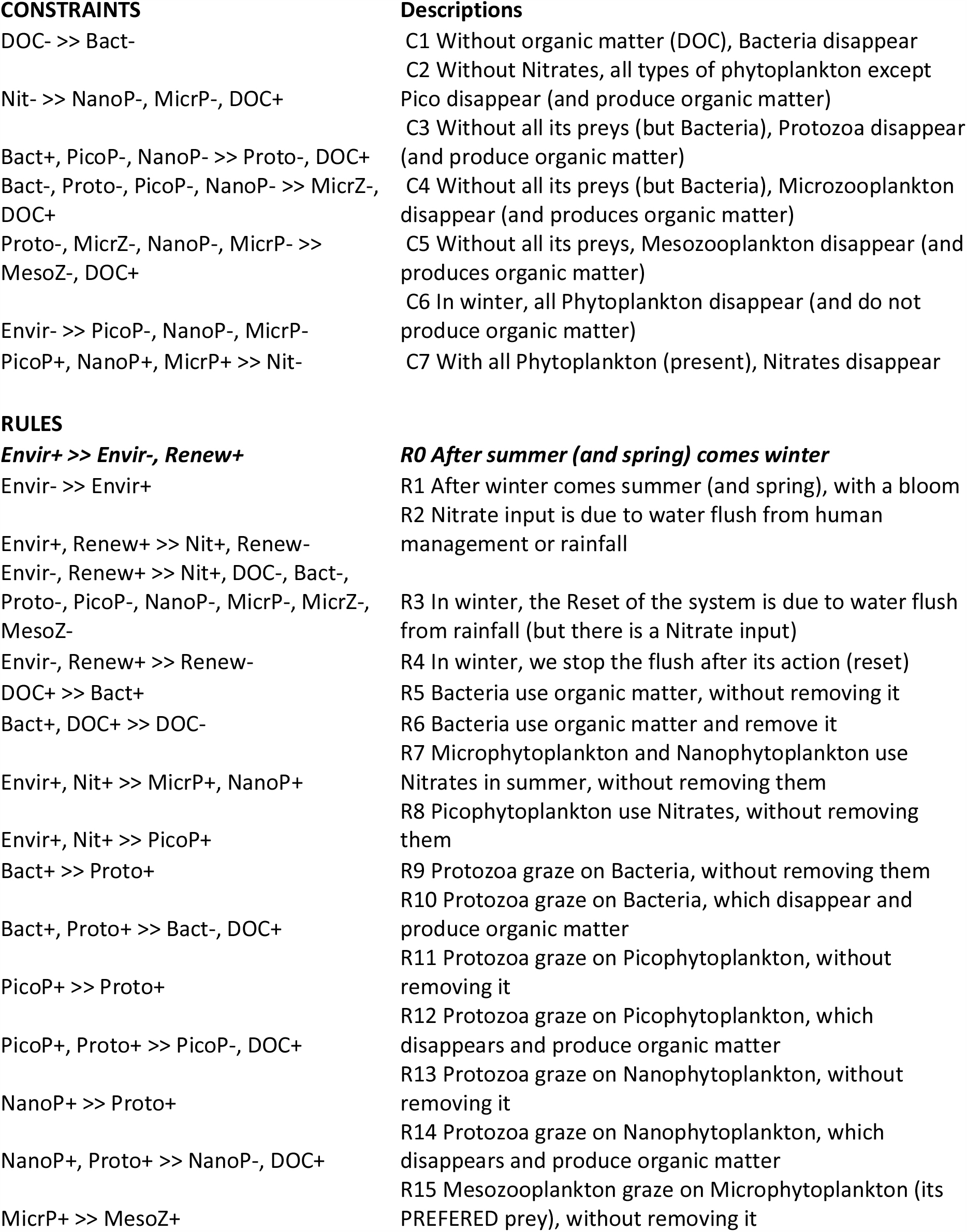
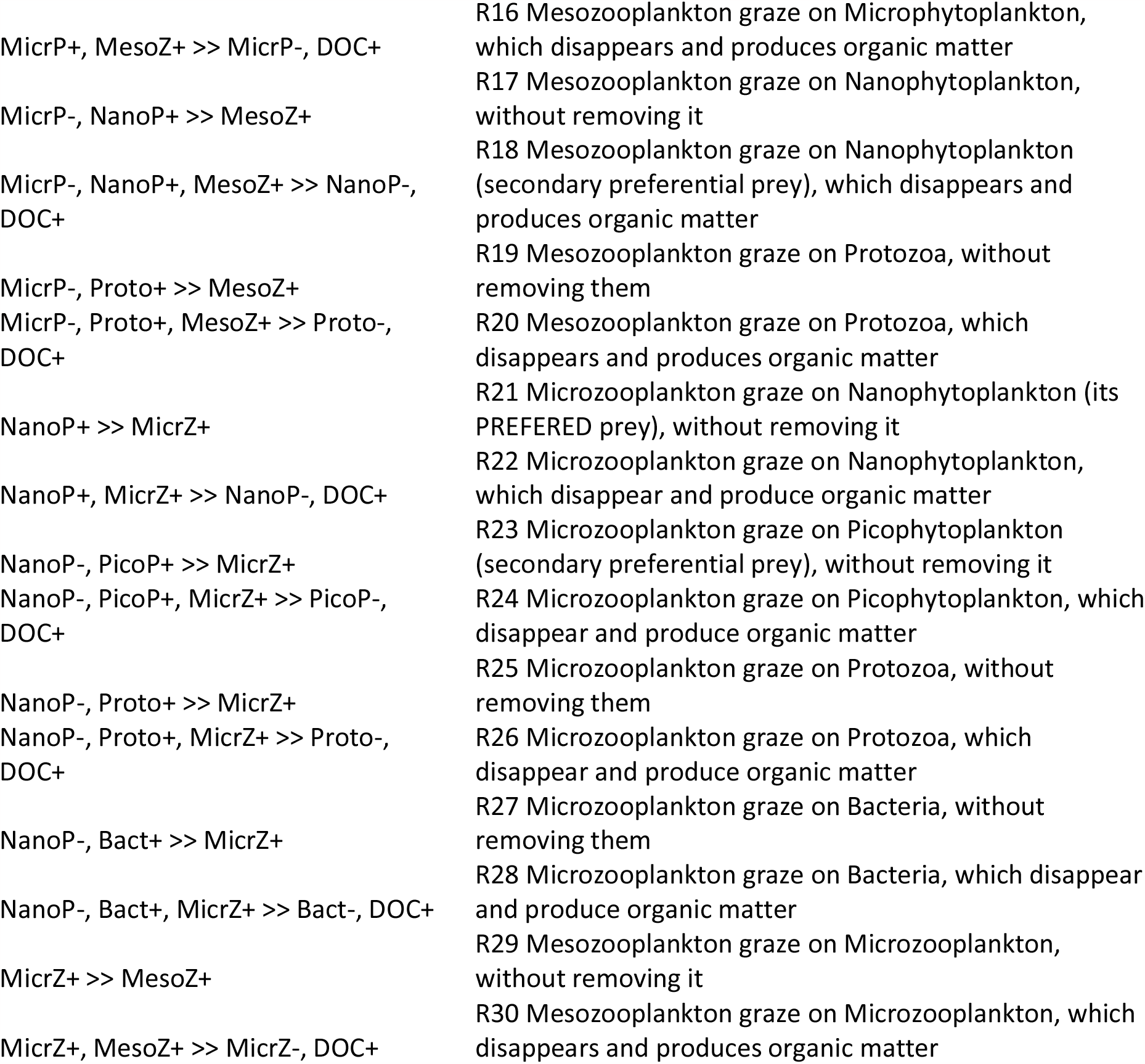
Rules and constraints used in both the full and seasonal models, with their formal expression (first column) and descriptions (second column). Only the rule N°0 (in italics and bold) is discarded in the seasonal version of the model.

For the full and seasonal models, we computed the state space and the *merged* state space, and we checked whether the observed trajectories were correctly recovered. A merged state space is a simplified state space gathering the sets of mutually reachable states of the modeled system, a topological structure called a *structural stability*, into the same nodes. This type of structure is interpreted as a stable regime as, by definition, any state in this stability can be reached by any other state belonging to it. Other stabilities can be identified automatically, such as *terminal stabilities*, from which the system can no longer exit, *basins* gathering states that have the same following states, and *deadlocks*, which are single states from which the system can no longer exit. Merged state spaces are much more compact than full state spaces, and summarized trajectories are readily revealed.

## RESULTS

### State spaces and computed dynamics

The full model provides a state space comprising 765 states gathered into a single dynamic structure (a so-called structural stability, Fig. SM1a). The seasonal model becomes stuck in a high number (12, plus two basins) of successive structural stabilities of various sizes (Figs. 4 and SM1b). When oriented downward in time; i.e., following causality and chronology, the whole system inevitably converges toward a small terminal stability (made up of four states) in which the system is in a biological winter (i.e., few living species, in green, Fig. SM1b). Some of the stabilities that are reached exhibit a large number of states and may keep the system into such specific stabilities for an indefinite time (in purple, Figs. 4 and SM1b). In brief, the seasonal model displays the same state space as the full model, but with a possible return to the initial biological winter regime, thus connecting the bottom states (in red, Fig. 4a) to the top states (in pink, Fig. 4a). This is why we observed a single cycling stability in the full model state space (Fig. SM1a).

**Figure 4.**
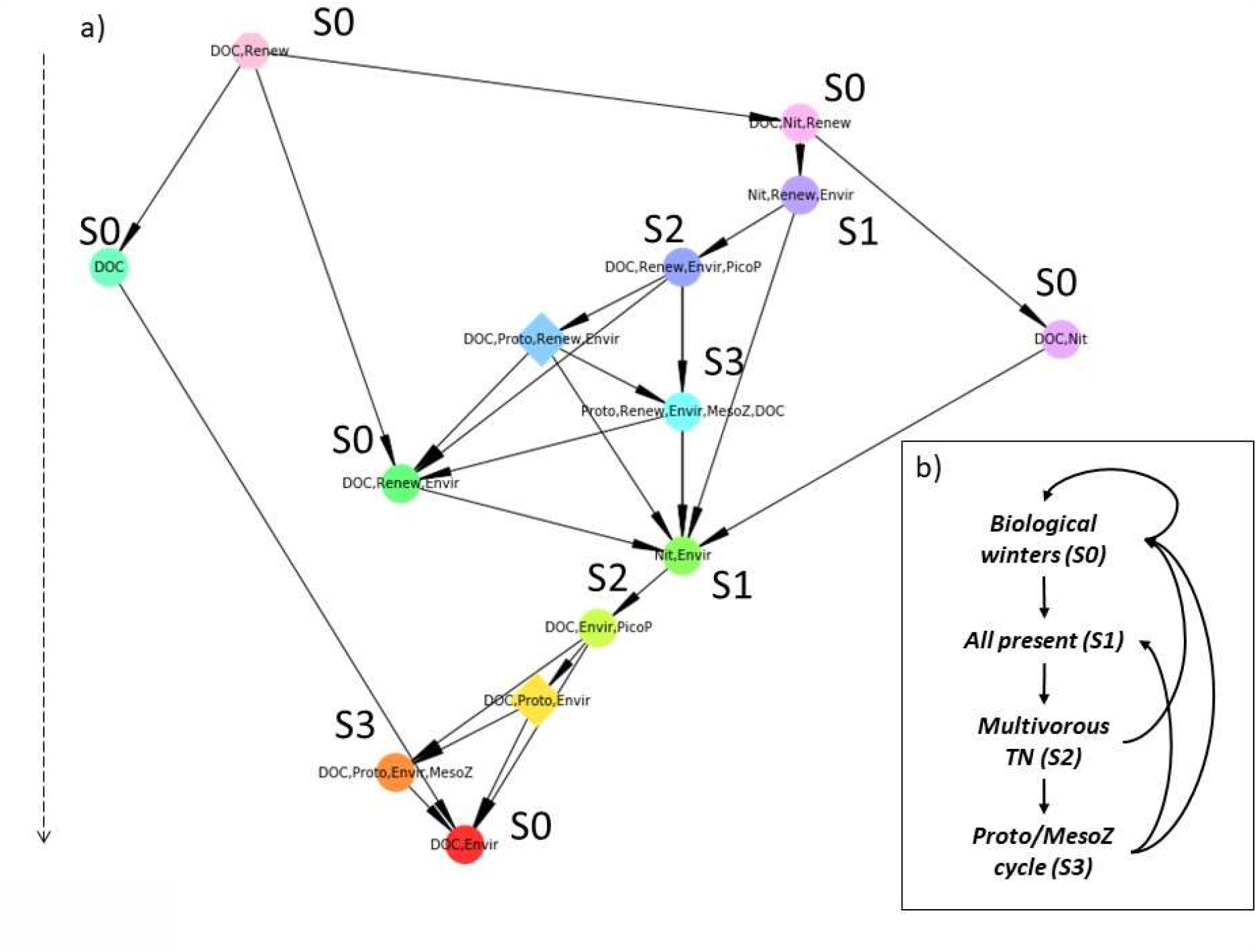
The merged state space of the seasonal model (a, as in Fig. SM1a), in which each node corresponds to a structural stability (i.e., a set of mutually reachable states), and each edge corresponds to irreversible transitions between successive stabilities. Here, structural stabilities are labeled with system components that are systematically present (+) in their associated states (see Fig. SM2). To see components that are systematically absent in stabilities). This figure helps identify the various regimes (b, and Table SM1) reached by the TN system along to the (downward) trajectories computed.

**Figure 5.**
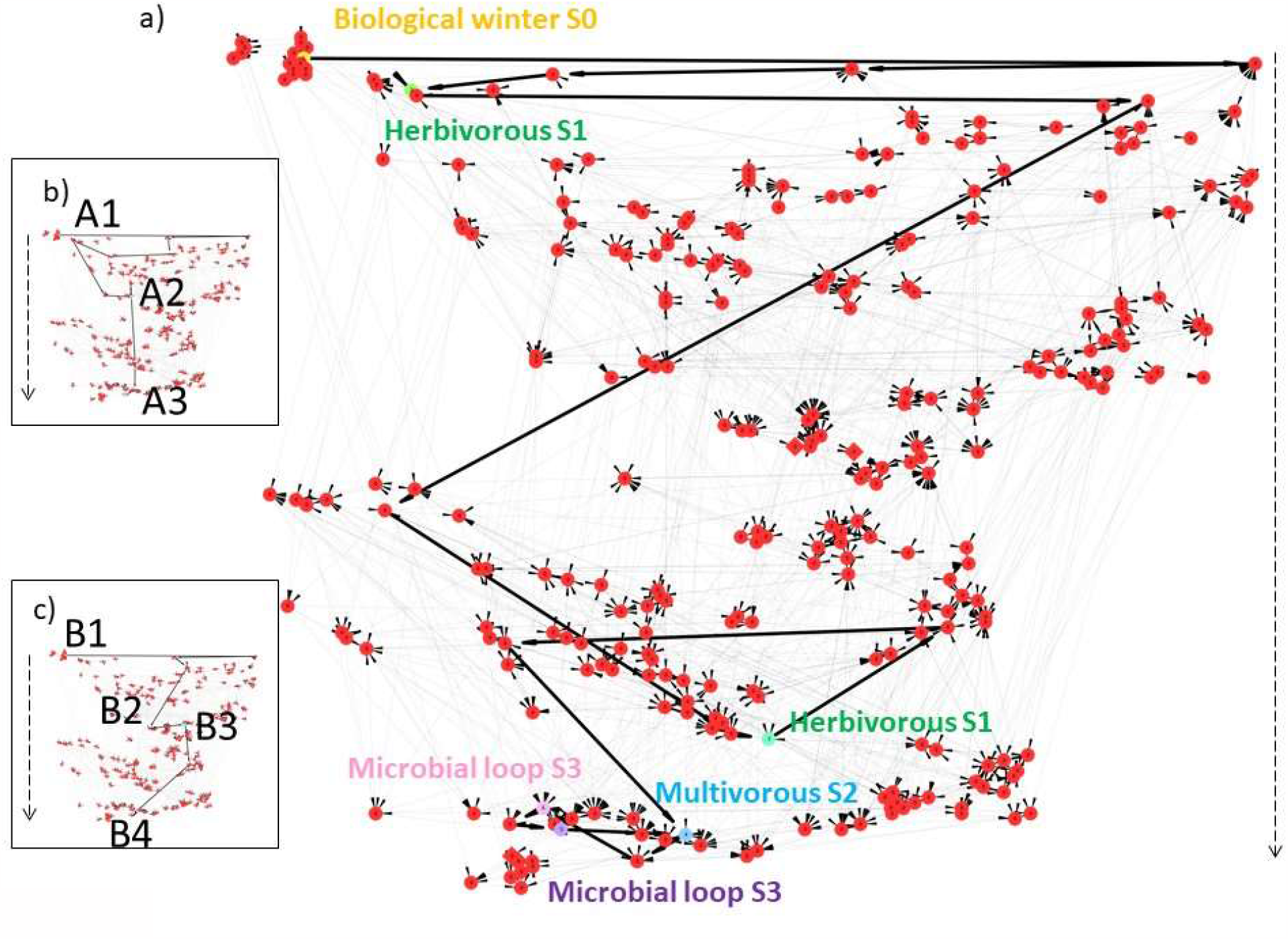
The full (not merged) seasonal state space highlighting the three trajectories used in this study (Table 2), namely the theoretical (a), station A (b), and station B (c) ones (Table 2 Suppl.). Here, each node corresponds to a TN state, connected to each other by downward transitions. The specific states underlying the three trajectories are highlighted by node colors other than red and identifiers corresponding to their numbers (last rows of trajectories in Table 2) and by bold edges.

Four TN regimes were revealed by the full and seasonal models (Fig. 4a and Table SM1): the S0 biological winter regime, without planktonic bloom, with oscillations of both zooplankton feeding on bacteria and organic matter; the S1 regime, in which all components are present because the environment is favorable to the development of organisms with many fluctuations of them; the S2 multivorous regime, with a mix of preys and various predators such as protozoa and both zooplankton (MicrZ and MesoZ), and finally, the S3 regime, centered on protozoa and mesozooplankton with a few preys but oscillation of Bacteria (Fig. 4a and Table SM1). The trajectories passing through different regimes were diverse and depended on the rules fired from the initial state (Fig. 4b): either the TN system shifts between various types of S0 regimes, or it successively crosses regimes S0 to S1, S2, and S3 (with possible ways back) and then back to S0.

### Model validation

All states of the theoretical trajectory were correctly predicted by the model and, as the model exhibited a single stability, the system is certain to successively reach all these states (although we do not know by which complicated trajectories, Fig. 4a, Table SM1). This observation definitely validates the model. The theoretical planktonic TN trajectory (Table 2 and Fig. 5a) started by an (immature) regime found during the biological winter. It then evolved toward low herbivorous TN, herbivorous TN, and variable multivorous TN (from weak to highly multivorous regimes, with protozoa, microzooplankton, or mesozooplankton, respectively), and a microbial TN regime, to finally reach a microbial loop regime. When the water in the marshes was renewed (*Renew*+), the TN returned to the biological winter regime, as can the herbivorous regimes as well. All these TN regimes were recovered by the model (Fig. 5a), yet with slightly different stabilities. Figure 5 is not meant to demonstrate this validation stage (already achieved by identification of the successive trajectory states), rather than *showing* that all predicted states (colored nodes) were correctly recovered in the computed state space, and indeed connected through successive transitions (bold edges). Note that this modeled trajectory crossed many other intermediate states (Fig. 5a) not found in the observations of (Masclaux et al. 2014). In the seasonal model (rule R0 deactivated), the theoretical trajectory was also predicted (colored states, Fig. 5a), yet with the last two states (blue states at the bottom) in the reverse order, as the fourth multivorous regime can directly reach the last depleted one.

The data recorders at stations A and B, and associated with observed trajectories, were also correctly recovered. At station A, three regimes succeeded over time, from A1 to A3 (Table SM2, and Fig. 5b). The TN started with biological winter for four weeks with the presence of nitrates and organic matter, but it did not reach favorable conditions for biological development. Then, the favorable conditions at week five allowed the development of phytoplankton (micro- and pico-plankton), and thus their zooplankton predators with bacteria. This situation was typical of situations between herbivorous and multivorous TNs. A multivorous regime of TN then took hold until week eight. At station B, the observed succession displayed four regimes, from B1 to B4 (Table SM2 and Fig. 5c). The TN started with biological winter for three weeks and favorable conditions occurring at week four, which allowed the presence of multivorous TN (“weak multivorous TN” according to (Masclaux et al. 2014)) and at week five an herbivorous TN. Then, a multivorous regime of TN took hold from week six to week eight.

## DISCUSSION

The discrete-event and qualitative model of trophic networks (TN) presented here can be computed instantaneously (< 0.01 s) and provided, once the model was defined and assumed, all possible trajectories of this system (Fig. 4). To our knowledge, this is the first attempt to exhaustively model a detailed TN (11 components, Table 1) and to accurately validate its qualitative dynamics.

### Complex dynamics of aquatic trophic networks

In the Charente-Maritime trophic system, we discovered that this TN may have followed other trajectories than the one identified by experts in the theoretical model and in the ones observed (Fig. 5 and Supplementary Materials tables). First, station B showed that *DOC* may be present in winter, thus with the TN fluctuating in intermediary states before reaching the usual trajectory observed in Masclaux et al. (2014). Indeed, *DOC* in winter could be an allochthonous input from the terrestrial environment (Del Gorgio and Davis 2003). After winter (i.e., when *Renew*+ and *Envir*+ were present, Table 1), all the modeled trajectories and all the TN regimes appeared at reach. The TN can return to biological winter system states due to the nitrate inputs (*Nit*+, with R2) and to anthropogenic activities (Tortajada et al. 2011). This situation occurs when the water renewal is substantial and no planktonic biomass accumulation is possible (David et al. 2020). Moreover, rainfall could occur and favor nitrate leaching (R3), thus pushing back the planktonic TN to biological winter system states. The model confirmed the key role of organic matter (*DOC*), as the system trajectories differed depending on whether or not organic matter was present at the beginning of winter.

From the initial state of the TN, the trajectories could pass through slightly different biological winter regimes (Figs. 4a-b) with oscillations in organic matter, bacteria, and micro- and mesozooplankton. Similarly, Masclaux et al. (2014) found two types of Biological winter regimes, mainly depending on the presence or absence of bacteria, and on some prey and predator combinations. The model correctly recovered different states of biological winters. The regime of multivorous TN was also well recovered by the model (Masclaux et al. 2014). The multivorous TN is known to be highly stable (Legendre and Rassoulzadegan 1995). However, the microbial loop, which has a transient nature (Legendre and Rassoulzadegan 1995) did not appear as a structural stability in the model either.

The regime gathering protozoa and mesozooplankton (*Proto*/*MesoZ* cycles) characterized by the presence of predators with a few preys but oscillation of bacteria was not found in the observations (Masclaux et al. 2014). The modeled trajectory crossed many intermediate states (Figs. 5a-c) not sampled in the field. The field sampling frequency or the structural characteristics of the sampled wetlands likely did not allow capture of all the possible states of TN: this reveals the ability of the model to explore many other possible states of the planktonic TNs and other trajectories of TN. In particular, the predicted *Proto*/*MesoZ* regime has not yet been identified at the Charente Maritime sites, but work in progress at other Atlantic arc territories has identified related TNs (F.-X. Robin, pers. comm.). Finally, bacteria were frequently present in the ecosystem, and they occupied a prominent place in the model (Table SM1–2, Table 2). Bacteria appeared to oscillate frequently (Fig. 4a), although this was not visible in the merged state space (i.e., bacteria frequently appear and disappear within structural stabilities). The model confirmed that bacteria are frequently grazed by their grazers, as are small protists (Pernthaler 2005, Šimek et al. 2013). Indeed, the high level of control of bacteria by the protozoa in freshwater ecosystems is already known.

### Power and drawbacks of discrete-event models

An increasing number of TN models have been developed (Mitra et al. 2007, Kriest et al. 2010, Thébault and Fontaine 2010, Turner et al. 2014, Kéfi et al. 2016, Hansen and Visser 2016, Kloosterman et al. 2016). But they still suffer from three main limitations: limited size and complexity, and a frozen (static) network with frozen (i.e., topology) interactions. In this study, we proposed a novel model family (called the EDEN framework (Gaucherel and Pommereau 2019, Cosme et al. 2022)) to bypass these limitations. Our model is based on a discrete-event system, well-known to computer scientists and more recently also some molecular biologists (Thomas and Kaufman 2001, Reisig 2013). The price to pay for using our qualitative model is that no quantitative and detailed dynamics are available; but in turn, no difficult parameterization and construction are required. Consequently, such an approach is fully complementary to already existing models in (trophic) ecology. Here, to the best of our knowledge, we provide for the first time a discrete and qualitative model of TN to bypass such limitations. Of note, in continuity with previous theoretical attempts (May 1973, Dambacher et al. 2003), we here open a new avenue for the use of such novel qualitative models in (ecosystem and trophic) ecology. The foundations of this proposition, yet beyond the present scope, are based on a theoretical consideration that assumes that ecosystems are informational systems comprised of material components and immaterial processes (Gaucherel 2019) summarized into their interaction networks.

Such a model is intuitive, easy to build, tractable, and rigorous (i.e., no trajectories have been forgotten or added according to the mathematical Petri net engine). In addition, we claim that it does not require any detailed or quantitative calibration, as no parameter is required. The central assumption of this approach is that it is possible to summarize ecological processes into qualitative rules, possibly interpreted as long-term and discrete events. Other studies have shown that this approach is not limited to trophic processes and can be applied to a high diversity of social-ecosystems (Gaucherel and Pommereau 2019, Mao et al. 2021). In this study, we were fortunate enough to collate several theoretical and observed trajectories with which to validate the model, thus confirming that it is conform and accurate (Fig. 5). Another quality of this type of model is to be heuristic, to force scientists questioning the knowledge they have regarding the studied system and to collate it into a single coherent framework.

As perspectives, it appears suitable to model many TN stressors such as pollution, cleaning, drought, invasive species, and/or climate changes (Mooney and Hobbs 2001, Mouquet et al. 2015). Any complexification of the studied social-ecosystem is also possible, in theory, as the model is still far from reaching its limits in terms of components, processes, and their nature diversity. It may then be used in a more applied manner, for exploration of other scenarios by changing the initial conditions. Coupling this model with other components describing the mechanisms behind these stressors would provide a relevant territorialized model to anticipate trends in a context of global warming and coastline changes. In the near future, it would be relevant not only to improve the model’s realism but also to develop analysis tools already used in similar studies focusing on social-ecological systems (Mao et al. 2021, Cosme et al. 2022). Additionally, it would be relevant to complexify our discrete and qualitative approach by using quantitative and multivalued schemes, to bridge the gap with more traditional (e.g., equation-based or individual-based) models (Vézina and Platt 1988, Kéfi et al. 2016).

In brief, by modeling trophic networks with a novel (EDEN) framework, we recovered theoretical as well as observed trajectories. With such qualitative models, the dynamics and predicted new states and new trophic network functioning regimes that may be observed in the field can be better understood. We illustrated these with a specific and well-documented freshwater trophic network. Such models provide an intuitive and robust approach to diagnosing any trophic (and non-trophic) network by computing all possible trajectories it can reach from a given chosen initial state. The known processes at play in the system help identify all of the possible dynamics and thus counter-intuitive trajectories of such complex (social-eco-)systems. Connecting such biotic dynamics to human-related activities can be expected to provide additional insightful understanding of trophic systems.

## Supporting information

Suppl Mat. Gaucherel et al. Aquatic Trophic

## APPENDICES

Additional Tables and Figures (Appendix 1)

## ACKNOWLEDGMENTS

We thank François-Xavier Robin for his relevant comments on an earlier draft of this paper.

## DATA SCRIPTS CODE AND SUPPLEMENTARY MATERIAL AVAILABILITY

Data are available in this article (Tables and Figures) (*citation of the data* Gaucherel et al, 2023);

Scripts and code are available online: DOI:10.1111/2041-210X.13242 of the webpage hosting the data https://github.com/fpom/ecco (*citation of the scripts and code* Pommereau et al., 2022);

Supplementary information is available online: XXXXDOI of the webpage hosting the data https://doi.org/10.5802/fake3.doi (*citation of the scripts and code* Gaucherel et al, 2023);

*[The references of the datasets, scripts and codes should also be present in the reference list and cited in the text*.]

