## Supplementary material for "Diagnosis of planktonic trophic network dynamics with sharp qualitative changes": Suppl Mat. Gaucherel et al. Aquatic Trophic

### Appendices

#### Additional Tables and Figures

**Table 1 Suppl.**: List of TN regimes computed by the model and found in the final state space (in Fig. 5). For each regime (columns), the present (+), absent (-) or oscillating (±) components of the trophic network are listed (in lines). This table helps identifying the various TN regimes crossed along to the computed trajectories (in Fig. 5 and 7).

| **State space identifier** | **S0** | **S1** | **S2** | **S3** |
|---|---|---|---|---|
| Bact | ± | ± | ± | ± |
| PicoP | - | + | + | - |
| NanoP | - | + | - | - |
| MicrP | - | ± | - | - |
| Proto | - | ± | ± | + |
| MicrZ | ± | ± | ± | ± |
| MesoZ | ± | ± | ± | + |
| DOC | ± | ± | ± | ± |
| Nit | ± | ± | - | - |
| Envir | - | + | + | + |
| Renew | + | - | - | - |
| **Regime names (Fig. 5 & 7)** | Biological winters | All component present | Multivorous TN | Proto/MesoZ cycle |

**Table 2 Suppl.**: Trajectories of the observed at stations A and B. For each trajectory, observed regimes are listed in columns and present (+)/absent (-) components of the TN are listed in lines. The corresponding regimes displayed in Fig. 6a-c are listed in the last line of each trajectory, with a single index A1 to A3 and B1 to B4 for successive regimes.

| **STATION A** | Week 1 | Week 2 | Week 3 | Week 4 | Week 5 | Week 6 | Week 7 | Week 8 |
|---|---|---|---|---|---|---|---|---|
| Bact | - | - | - | - | + | + | + | + |
| PicoP | - | - | - | - | + | + | + | + |
| NanoP | - | - | - | - | - | - | - | - |
| MicrP | - | - | - | - | + | + | + | + |
| Proto | - | - | - | - | - | + | + | + |
| MicrZ | - | - | - | - | + | - | - | - |
| MesoZ | - | - | - | - | + | + | + | + |
| Nit | + | + | + | + | + | - | - | - |
| DOC | - | - | - | - | + | + | + | + |
| Envir | - | - | - | - | + | + | + | + |
| Renew | + | + | + | + | + | - | - | - |
| **Regimes (Fig. 6)** | **A1** | **A1** | **A1** | **A1** | **A2** | **A3** | **A3** | **A3** |

| **STATION B** | Week 1 | Week 2 | Week 3 | Week 4 | Week 5 | Week 6 | Week 7 | Week 8 |
|---|---|---|---|---|---|---|---|---|
| Bact | - | - | - | + | + | + | + | + |
| PicoP | - | - | - | + | + | + | + | + |
| NanoP | - | - | - | - | - | + | + | + |
| MicrP | - | - | - | + | + | - | - | - |
| Proto | - | - | - | + | - | - | - | - |
| MicrZ | - | - | - | - | + | + | + | + |
| MesoZ | - | - | - | + | + | + | + | + |
| Nit | + | + | + | - | + | - | - | - |
| DOC | + | + | + | + | + | + | + | + |
| Envir | - | - | - | + | + | + | + | + |
| Renew | + | + | + | + | + | - | - | - |
| **Regimes (Fig. 6)** | **B1** | **B1** | **B1** | **B2** | **B3** | **B4** | **B4** | **B4** |

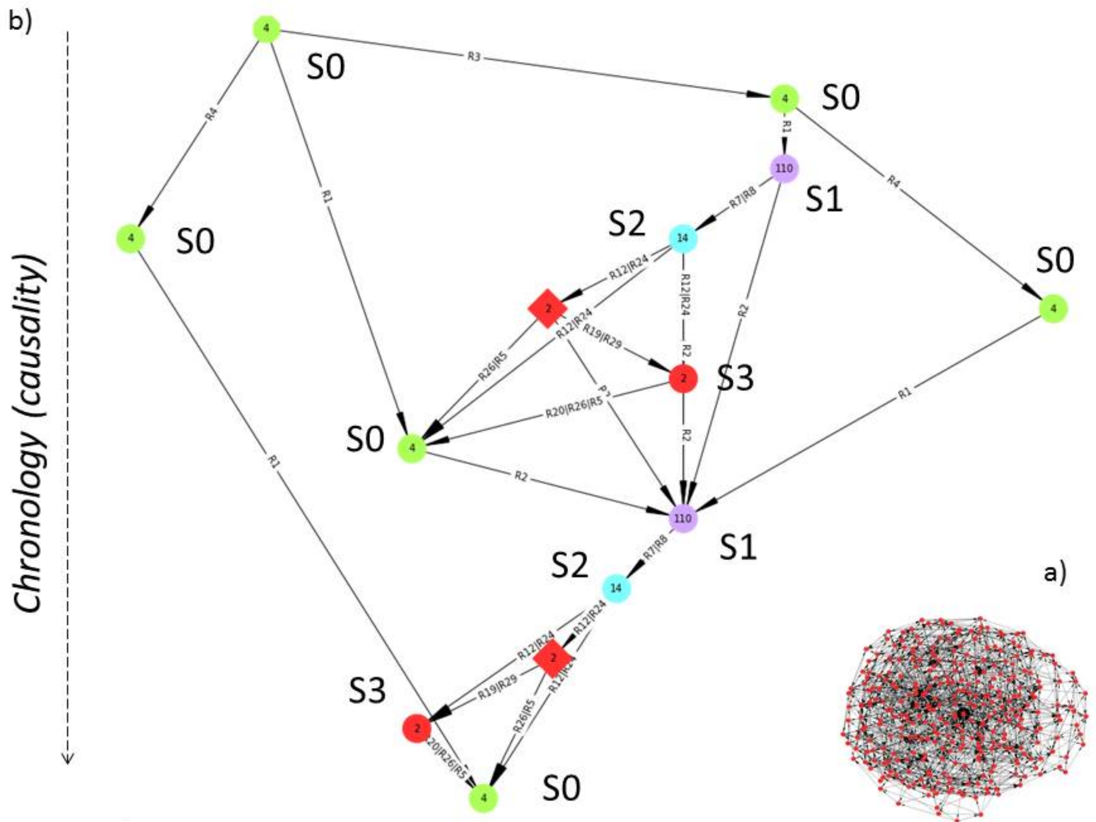


Figure 1 Suppl.

**Figure 1 Suppl.** The merged state spaces of the full (a) and seasonal (b) models, on which each node corresponds to a structural stability (i.e. a set of TN states), and each edge corresponds to specific transitions between two successive stabilities. The seasonal merged state space should be read from top to bottom (following causality), through the various trajectories connecting each structural stability and TN regimes (see main text). The structural stabilities of various types (node shapes, round shape for stabilities and lozenge for basins) are labeled by the number of states they are composed of (with distinctive colors), while the transitions between them are labeled by the rules allowing the changes.

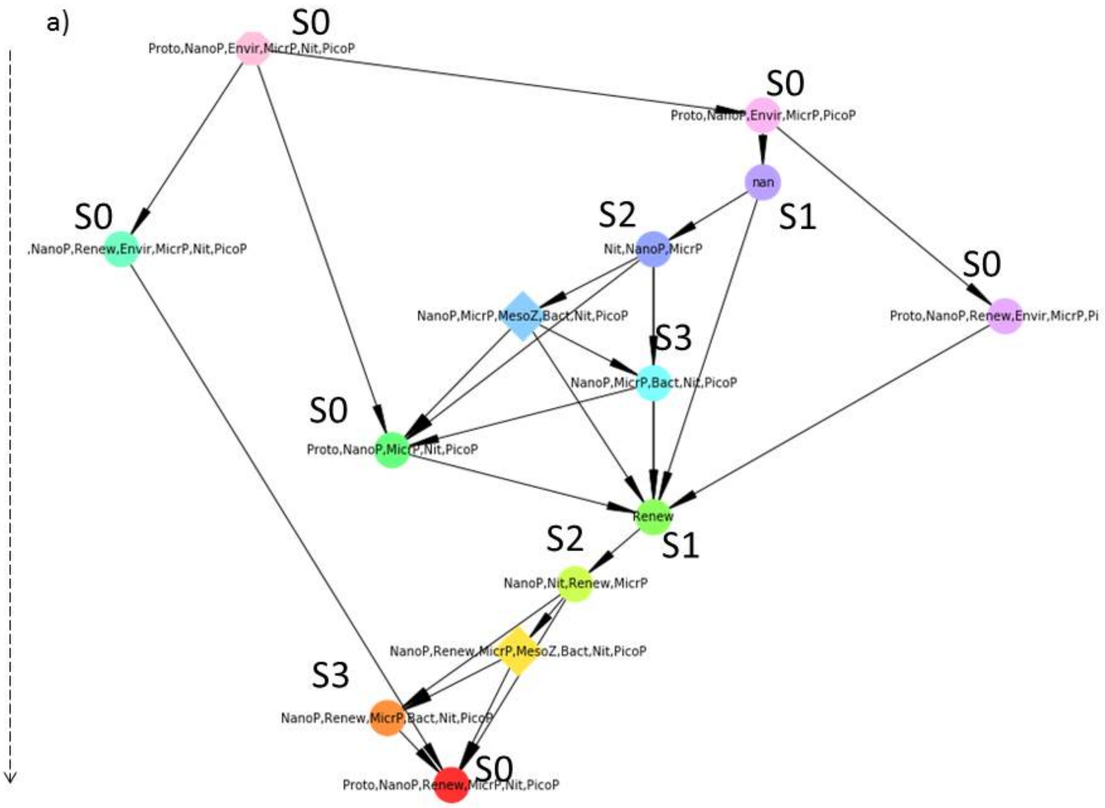


Figure 2 Suppl.

**Figure 2 Suppl.**: The same seasonal merged state space (a) as in Fig. 5a and 1 Suppl., yet with structural stabilities labeled with system components that are systematically absent (-) in their associated states. This figure helps identifying the various regimes (b, table 2 Suppl.) reached by the TN along to the (downward) trajectories computed. The TN components that are not systematically present (Fig. 5a) nor systematically absent (this figure) are oscillating (noted ± in the table 2 suppl.).
